# Individual neurons in monkey sensory and prefrontal cortices contribute to both working memory maintenance and readout

**DOI:** 10.64898/2026.03.13.711560

**Authors:** Ying Huang, Michael Brosch

**Author notes:** Correspondence: Ying Huang, PhD. Author contributions: Conception and Design: Y.H. and M. B; Data Acquisition and Analysis: Y.H.; Data Interpretation: Y.H. and M.B; Writing: Y.H. and M.B. Competing interests: The authors declare that they have no competing interests.

## Abstract

Working memory enables the use of past sensory information to guide behavior, yet how maintained information is linked to behavioral readout remains unclear. Using a dual-task paradigm to isolate working memory from concurrent mental processes, we demonstrate that the same neurons in the auditory and prefrontal cortices of monkeys maintain stimulus-specific information across a delay and are subsequently engaged in reading out that information in a manner reflecting the animal’s behavioral decisions. The functional relevance of these neurons is supported by causal perturbations of the dopaminergic system, which selectively impair working memory performance while leaving other task-related processes intact. Crucially, the maintenance-readout coupling is obscured under traditional single-task approaches that rely on comparisons of delay and baseline periods. The presence of this coupling in both the auditory and prefrontal cortices suggests a conserved computational architecture across the sensory–prefrontal axis, rather than a principle exclusive to executive centers. Together, these results demonstrate that memory maintenance and behavioral readout are computationally integrated, suggesting a unified neuronal framework for bridging past sensory input with future action.

**Significance Statement:** Working memory allows us to maintain information over short periods and use it to guide behavior. It is often assumed that these two functions, maintenance and readout, are carried out by separate neuronal populations. Here, we show that the same neurons in sensory and prefrontal cortices of monkeys both maintain stimulus-specific information and contribute to behavioral decisions. Using a dual-task paradigm that dissociates working memory from other cognitive processes, we identify neuronal roles that would be obscured under traditional single-task designs. These findings provide a unified framework for studying working memory and reveal how memory representations can be directly translated into actions within the same neuronal circuits.

## 1. <u>Introduction</u>

Working memory (WM) is the mental process that temporarily maintains information online so that it can be read out at the appropriate time to guide goal-directed behavior^1–4^. Studies investigating the neural implementation of WM typically employ tasks with a delay period following a sample stimulus, culminating in a test stimulus that prompts a behavioral output. These studies have identified neural activity during the memory-demanding delay that is selective for specific stimuli. Such delay activity occurs in multiple cortical regions, from sensory areas to the prefrontal cortex (PFC), and is conventionally regarded as the neural substrate of WM maintenance^5–12^. However, it remains unclear how these memory representations are read out and translated into behavior, specifically, whether this transformation occurs directly within the maintenance circuits or requires relaying the memory to other neuronal populations.

Addressing this issue is complicated by profound analytical challenges related to characterizing the precise functional role of delay activity within the maintenance circuits. Traditionally, delay activity is identified by contrasting memory periods with baselines recorded before stimulus presentation or task engagement. While foundational for mapping task-related modulation, this delay-baseline comparison inherently captures a mixture of multiple neural phenomena. For example, even without task engagement, stimulus-evoked activity can naturally outlast stimulus presentation and remain stimulus-selective, a well-documented feature of both sensory and prefrontal cortices^13–16^. Consequently, disentangling residual sensory processing from memory maintenance presents a substantial analytical difficulty for traditional delay-baseline comparisons.

Moreover, when subjects perform WM tasks, the memory delay involves several concurrent mental processes required for task performance, such as reward or stimulus anticipation, decision making, and action preparation. Neural signals tuned to these task variables often persist for several seconds^17–25^. Therefore, delay activity identified through delay-baseline comparisons within a single task can reflect these variables alongside WM itself. This mixing of neural signals may explain why previous studies have reported at most a weak relationship between delay activity and the neural signatures observed during the subsequent memory readout phase^26–32^. This apparent dissociation has fueled ongoing debates over the functional architecture of WM, namely, whether memory maintenance and readout are handled by anatomically segregated neuron populations or are functionally integrated within common circuits^2,11,33–37^. This leaves a critical gap in our understanding of how information maintained in WM is transformed into a motor action.

To isolate these concurrent variables, dual-task designs have been developed to compare delay periods across tasks with different WM demands^38–40^. In these implementations, however, WM demands often co-varied with other factors, such as using stimuli from distinct sensory modalities^38^ or varying stimulus durations^39^. The delay activity identified under these conditions may partially reflect differences in stimulus-evoked activity that either sustains throughout or outlasts the stimulus presentation^13–15,41,42^. Even when identical stimuli were deployed in our previous work^40^, subsequent analyses revealed that subjects might have adopted behavioral strategies not involving WM^43,44^. This indicates that further paradigm refinement is required to definitively isolate the mnemonic components of delay activity.

We thus used a highly controlled dual-task paradigm in monkeys to directly examine the neural architecture that links the maintenance of WM information to subsequent behavioral readout. We trained monkeys on two tasks that differed in auditory WM demand during the delay but were otherwise identical in the acoustical stimulation prior to this period and in other task variables. We recorded spiking activity across the frontotemporal axis, targeting the auditory cortex (AC) and PFC, namely, the two keys nodes conventionally implicated in auditory WM^31,32,40,45^. By comparing these regions, we tested whether sensory and prefrontal cortices follow similar functional principles to maintain and read out memory representations. After characterizing this circuit organization, we tested for causality by perturbing neuronal activity via microinjections of dopaminergic drugs. We specifically targeted AC because the necessity of PFC for WM is already established, whereas the extent to which local dynamics within sensory cortices are required remains a major open question. We focused on the dopaminergic system, as it is a critical regulator of WM^46,47^ and modulates processing within AC^48,49^.

## 2. Results

### 2.1. Spiking activity in auditory cortex during the delay period is related to auditory working memory

Monkey C and Monkey L performed two tasks on sequences of 200-ms sounds S1 and S2, separated by an 800-ms silent delay (Fig. 1A). Cued by LED signals, they triggered sound presentation by grasping a touch bar. In the delayed match-to-sample (DMS) task, cued by red LEDs on the left, the monkeys reported whether S1 and S2 were identical or different, via a *Go* response (releasing the bar after S2 offset) or a *NoGo* response (continuing to hold it), respectively. Both S1 and S2 could be a 0.4-, 1.2-, or 3.6-kHz tone. Here, monkeys had to maintain auditory information in WM during the delay until S2, and then read it out, retrieving and relating it to S2, to decide on a Go or a NoGo response. The same three tones served as S1 in the S2-discrimination (S2D) task, cued by green LEDs on the right, where S2 could be a noise burst, a 20-Hz or an 80-Hz click train; a Go response was required after noise bursts, and NoGo otherwise. Since the required response was independent of S1, monkeys did not need to maintain auditory information in WM during the delay. Thus, the two tasks differed in auditory WM demand during the delay, while other task variables, such as decision making, action preparation, and procedural demands, were matched.

**Fig. 1.**
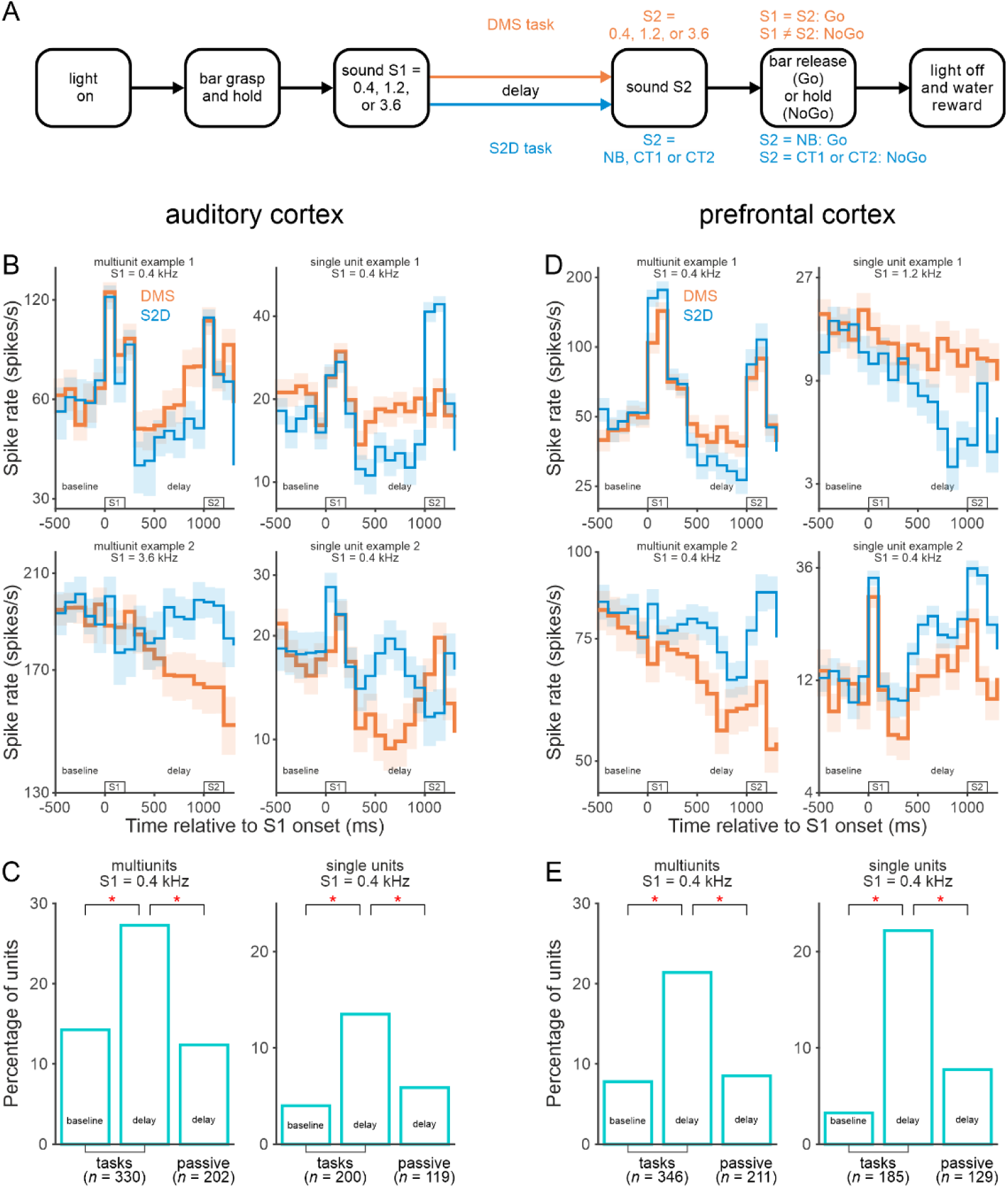
Delay activity related to auditory working memory in auditory and prefrontal cortices revealed by a dual-task approach. **(A)** Behavioral tasks. After LED onset, the monkeys had to grasp a bar and hold it, triggering a sequence of two sounds, S1 and S2, separated by a delay. In both the delayed match-to-sample (DMS) task (orange) and S2-discrimination (S2D) task (blue), S1 could be a 0.4-, 1.2- or 3.6-kHz tone. These tones were also used as S2 in the DMS task, in which monkeys had to make a Go response when S2 matched S1 and a NoGo response when it differed. In the S2D task, S2 could a noise burst (NB), a 20-Hz click train (CT1) or an 80-Hz click train (CT2). A Go response was required when S2 was a noise burst and a NoGo response otherwise. **(B)** Four example WM units in auditory cortex (AC) exhibiting significantly different spike rates during the final 500 ms of the delay between the DMS and S2D tasks. Spike rates of multiunits (MUs; left panels) and single units (SUs; right panels) were averaged across correct trials with a 0.4- or a 3.6-kHz S1. Shaded area indicates standard error of the mean. **(C)** Percentage of AC units displaying significant delay or baseline spike ratios. Ratios were calculated from trials with a 0.4-kHz S1, either between the two tasks or between the corresponding passive conditions (identical acoustic stimulation without task engagement). Delay spike ratios were computed from the final 500 ms of the delay; baseline spike ratios from the 500 ms preceding S1. The total sample size (*n*) for analyzed MUs and SUs are indicated below each condition. Asterisks denote significant differences between percentages. Note that significant ratios occurred most frequently during the delay period of the tasks. **(D, E)** WM-related delay activity observed in prefrontal cortex (PFC). Conventions for panel D are identical to those for panel B, with the PFC example units illustrated for the 0.4- or 1.2-kHz S1. Conventions for panel E are same as those for panel C.

The two tasks were performed in alternating blocks of trials in 134 sessions without pharmacological perturbation (72 from Monkey C and 62 from Monkey L). We recorded spiking activity from core fields of the right AC in 123 sessions and from the ventrolateral area of the left PFC in 115 sessions.

Focusing first on these non-pharmacological sessions, we leveraged the dual-task paradigm to isolate the mnemonic component of delay-period activity in AC. Numerous units in AC exhibited spiking activity related to auditory WM. Figure 1B illustrates how example multiunits (MUs) and single units (SUs) fired in correctly performed DMS (orange) or S2D trials (blue) with a 0.4-kHz S1 (top panels, bottom-right panel) or a 3.6-kHz S1 (bottom-left panel). All four units fired differently between the two tasks shortly after S1, with the difference persisting throughout the remainder of the delay. To quantify this, we computed for each unit the ratio of the mean spike rate during the final 500 ms of the delay between tasks (DMS / S2D). These delay spike ratios for the top-panel units, 1.40 for the MU (left) and 1.51 for the SU (right), were significantly greater than 1 (p<0.05, permutation test), indicating higher firing rates in the DMS task. In contrast, the bottom-panel units had significant delay spike ratios less than 1 (0.88, left; 0.63, right), reflecting lower firing rates in the DMS task.

From correct trials with a 0.4-kHz S1, significant delay spike ratios were found in 90 of all 330 MUs (27.3%; Fig. 1C left, middle bar) and in 27 of all 200 SUs (13.5%; Fig. 1C right, middle bar), with values either above or below 1 (MUs: 64 vs. 26; SUs: 15 vs. 12). We next demonstrate that these significant ratios genuinely reflected differences in auditory WM between the tasks rather than potential confounding factors.

Because S2 differed between tasks, the across-trial acoustic context also differed between DMS and S2D blocks (trials preceded by a tone vs. a complex sound). To assess effects of this acoustic context on neuronal activity^50^, we analyzed spiking activity during passive sound presentation, computing delay spike ratios between passive blocks containing the same sound sequences as the DMS and S2D tasks. This analysis revealed that the significant delay spike ratios could not be solely accounted for by differences in acoustic context between DMS and S2D blocks. For both MUs and SUs, significant delay spike ratios occurred in passive conditions, but significantly less often than during task performance (Fig. 1C, right vs. middle bars; MUs: 12.4% of 202 vs. 27.3% of 330; SUs: 5.9% of 119 vs. 13.5% of 200; p<0.05, permutation test).

The significant delay spike ratios during task performance could also not be simply explained by differences in light stimulation. We computed spike ratios between tasks during a period closer to light onset than the delay: the 500-ms baseline preceding S1. While some of these baseline spike ratios were significant, their proportions were significantly smaller than during the delay (Fig. 1C, left vs. middle bars; MUs: 14.2% vs. 27.3%; SUs: 4.0% vs. 13.5%; p<0.05, permutation test). This result indicates that the significant delay spike ratios cannot be accounted for by long-lasting effects of light stimulation on spiking activity^25^, which would predict equal or higher prevalence during baseline. It also suggests that these significant ratios cannot be attributed to differences in reward anticipation between tasks^20^. Such differences might arise from the higher accuracy in the S2D task (median: 87% vs. 70% in the DMS task; 134 sessions), leading to more frequent rewards and potentially stronger anticipation. However, these differences would have produced significant spike ratios at similar frequencies during the delay and baseline.

Based on these results, we concluded that the significant delay spike ratios observed in the 90 MUs and 27 SUs can be at least partially attributed to differences in auditory WM between tasks. Such WM-related delay activity was also obtained from sound sequences with S1s other than 0.4 kHz. Specifically, significant delay spike ratios were found in 88 MUs and 29 SUs after the 1.2-kHz S1, and in 86 MUs and 31 SUs after the 3.6-kHz S1.

Significant delay spike ratios after the 0.4-, 1.2-, or 3.6-kHz tone were detected both in units responding to that same tone in the tasks and those not responding to it (Fig. 2A; bars to the right and left of the vertical line, respectively; see Section 4.6 for unit classification; see Fig. 1B for examples). For MUs, these ratios occurred with comparable or higher probability in responsive than in non-responsive populations (27.6% vs. 29.1%, 30.4% vs. 21.6%, and 28.1% vs. 15.6% for 0.4, 1.2, and 3.6 kHz, respectively), with the difference being significant after the 3.6-kHz S1 (p<0.05, permutation test). For SUs, significant ratios occurred similarly often across the two populations (SUs: 10.6% vs. 16.0%, 16.3% vs. 14.9%, and 16.4% vs. 15.5%; p> 0.05 for all three comparisons). These observations suggest that both stimulus-encoding and non-encoding neurons are involved to maintain information during auditory WM tasks.

**Fig. 2.**
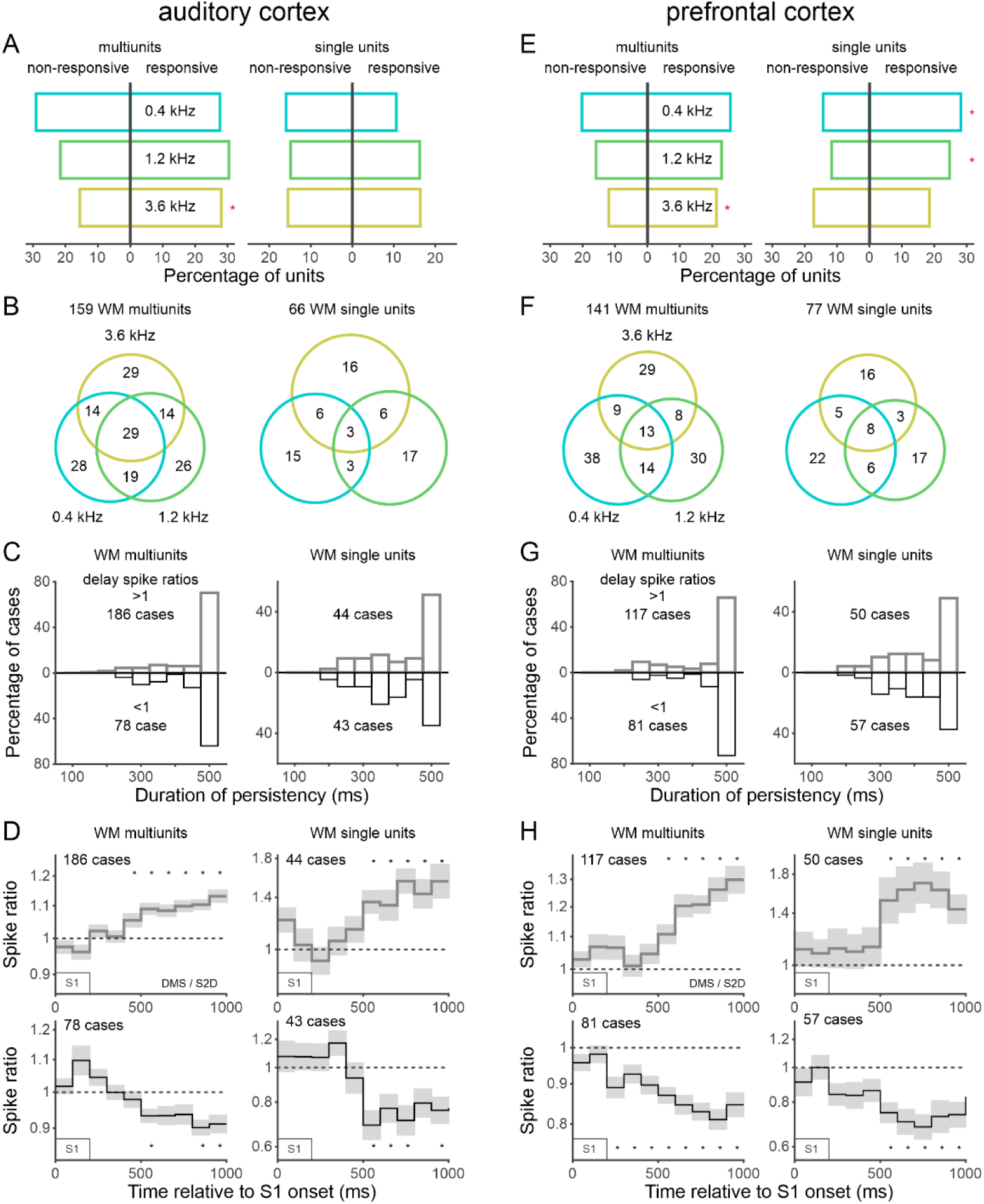
Characteristics of working memory-related delay activity in auditory and prefrontal cortices. **(A)** Both tone-responsive and non-responsive AC units exhibited WM-related delay activity. Unit classification was performed separately for the 0.4-, 1.2-, and 3.6-kHz tones, and delay spike ratios were assessed for the corresponding tone in each case. Bars to the right and left of the vertical line indicate the percentages of responsive and non-responsive units, respectively, with significant delay spike ratios. Significant differences between these percentages are marked by asterisks. Data obtained from MUs (left) and SUs (right) are shown separately. Sample sizes for the 0.4-, 1.2-, and 3.6-kHz tone, respectively—responsive MUs: 152, 101, and 90; non-responsive MUs: 141, 161, and 171; responsive SUs: 60, 80, and 55; non-responsive SUs: 119, 101, and 116. **(B)** Most AC units exhibited WM-related delay activity only for specific tones. The Venn diagrams illustrate distributions of units showing a significant delay spike ratio for the 0.4-, 1.2-, or 3.6-kHz tone. **(C)** Many AC MUs and SUs exhibited WM-related activity throughout the final 500 ms of the delay. Distributions show the cumulative duration of consecutive windows displaying spike ratios > 1 or <1 during the delay between the two tasks. Spike ratios were calculated in 100-ms sliding windows, with 50-ms steps, for individual cases where WM units exhibited a significant delay spike ratio for the 0.4-, 1.2-, or 3.6-kHz tone, plotted separately for cases >1 (gray bars; above the horizontal line) or <1 (black bars; below the horizontal line). Note that, in about 70% of MU cases and 40% of SU cases, the total duration reached the maximum value of 500 ms. **(D)** Ramping dynamics of AC population spike ratios calculated between the two tasks during consecutive 100-ms windows aligned to S1 onset. Data are shown separately for MU and SU cases with significant delay spike ratios >1 (top row) or <1 (bottom row). Population spike ratios continued to deviate from 1 after S1 offset, with the deviation being largest near the end of the delay. Asterisks mark significant spike ratios. **(E-H)** WM-related delay activity in PFC exhibit features common to those observed in AC. It is distributed across both tone-responsive and non-responsive sub-populations (E), is predominantly tone-specific (F), and persists throughout the delay, with ramping dynamics toward the end of the delay (G, H). In panel E, PFC sample sizes for the 0.4-, 1.2-, and 3.6-kHz tone, respectively—responsive MUs: 121, 127, and 141; non-responsive MUs: 188, 189, and 176; responsive SUs: 103, 85, and 65; non-responsive SUs: 77, 94, and 105.

The overall 264 significant MU and 87 significant SU delay spike ratios after the 0.4-, 1.2-, or 3.6-kHz tone were distributed across 159 MUs (Fig. 2B, left Venn; 48.2% of 330) and 66 SUs (right Venn; 33.0% of 200), respectively. These units were classified as WM units.

Most of the WM units showed significant delay spike ratios after only one tone (83 MUs and 48 SUs) or two tones (47 MUs and 15 SUs), while a minority did so following all three (29 MUs and 3 SUs; Fig. 2B). Thus, 81.8% of WM MUs and 95.5% of WM SUs exhibited memory-related activity that was tone specific (p<0.05, permutation test). This pattern was consistent across WM units responsive to the 0.4-, 1.2-, or 3.6-kHz tone (81.8% of 126 MUs; 97.6% of 41 SUs) and those responsive only to other sounds (81.8% of 33 MUs; 92.0% of 25 SUs). These results suggest that stimulus-specific information is maintained by both stimulus-encoding and non-encoding neurons.

Many WM units exhibited memory-related activity that persisted throughout the delay (see examples in Fig. 1B). We revealed this by calculating spike ratios between tasks on a finer time scale during the final 500 ms of the delay, using 100-ms sliding windows with 50-ms steps. The calculation was performed separately for sequences with different S1s and restricted to cases with significant delay spike ratios. These spike ratios remained consistently above or below 1 throughout the entire 500 ms in 68.6% of MUs and 43.7% of SUs (Fig. 2C, rightmost bar). In most remaining cases, ratios stayed consistent for ≥300 ms, namely, more than half of the entire 500 ms (MUs: 81.9%; SUs: 77.6%). These findings indicate that WM-related activity was predominantly persistent.

The persistence of WM-related activity was further examined by computing the time course of population spike ratios for the MU and SU cases, using consecutive 100-ms windows aligned to S1 onset (Fig. 2D). For cases with significant delay spike ratios >1, population values hovered around 1 shortly after S1 onset, became significantly larger than 1 about 400 ms after S1 offset, and peaked toward the end of the delay (top row; α=0.005, corrected across ten bins, permutation test). A mirrored ramping dynamic was observed for cases with significant delay spike ratios <1, with population values reaching a minimum near the end of the delay (bottom row). These dynamics indicate that WM-related activity became gradually more prominent during the delay, peaking just before S2, precisely when the information maintained in WM must be read out to guide behavior.

### 2.2. Working memory units in auditory cortex are involved in reading out maintained information

To determine whether these WM units, which belong to the maintenance circuit, are also involved in reading out maintained information to guide behavior, we analyzed their activity during the readout phase of the DMS task. This phase is when S2 was presented and had to be related to S1 to determine the appropriate action: releasing the bar (Go) if it matched S1 and withholding release (NoGo) if it did not. For each unit and tone (0.4-, 1.2-, and 3.6-kHz), we computed post-stimulus time histograms (PSTHs; 50-ms bins) aligned to S2 onset for match and non-match trials. This computation was restricted to cases where the unit exhibited significant delay spike ratios for that specific tone. PSTHs were normalized to the 100 ms before S2 and averaged across all cases to obtain population spike rates, separately for units responsive to the corresponding tone and for those not responsive.

Figure 3A shows population spike rates for tone-responsive WM MUs (left) and WM SUs (right) during correct task performance. Spike rates were generally lower in match than in non-match condition, with differences emerging shortly after S2 onset and lasting throughout its 200-ms presentation (red vs. black traces). These differences were significant from 50 to 150 ms after S2 onset for MUs and from 0 to 100 ms for SUs (α=0.005, corrected across ten post-S2 bins). Consistently, normalized mean spike rates across the 0-200 ms period were lower in the match condition for most MUs (68.3%) and SUs (70.0%; Fig. 3B, dots below diagonal), with the MU proportion significantly exceeding 50% chance (p<0.05, permutation test). This match-related reduction occurred both in units with delay spike ratios >1 (grey dots; MUs: 72.1%; SUs: 75.0%) and those with ratios <1 (black dots; MUs: 59.5%; SUs: 66.7%). Thus, regardless of how these units maintained information during delay, they generally exhibited reduced activity during the readout phase when S2 matched S1. Subsequent analyses suggest that these activity differences reflected the monkeys’ match judgement and Go decision, rather than motor execution of bar release or stronger forward inhibition in match trials where S1 and S2 shared the same frequency^50^.

**Fig. 3.**
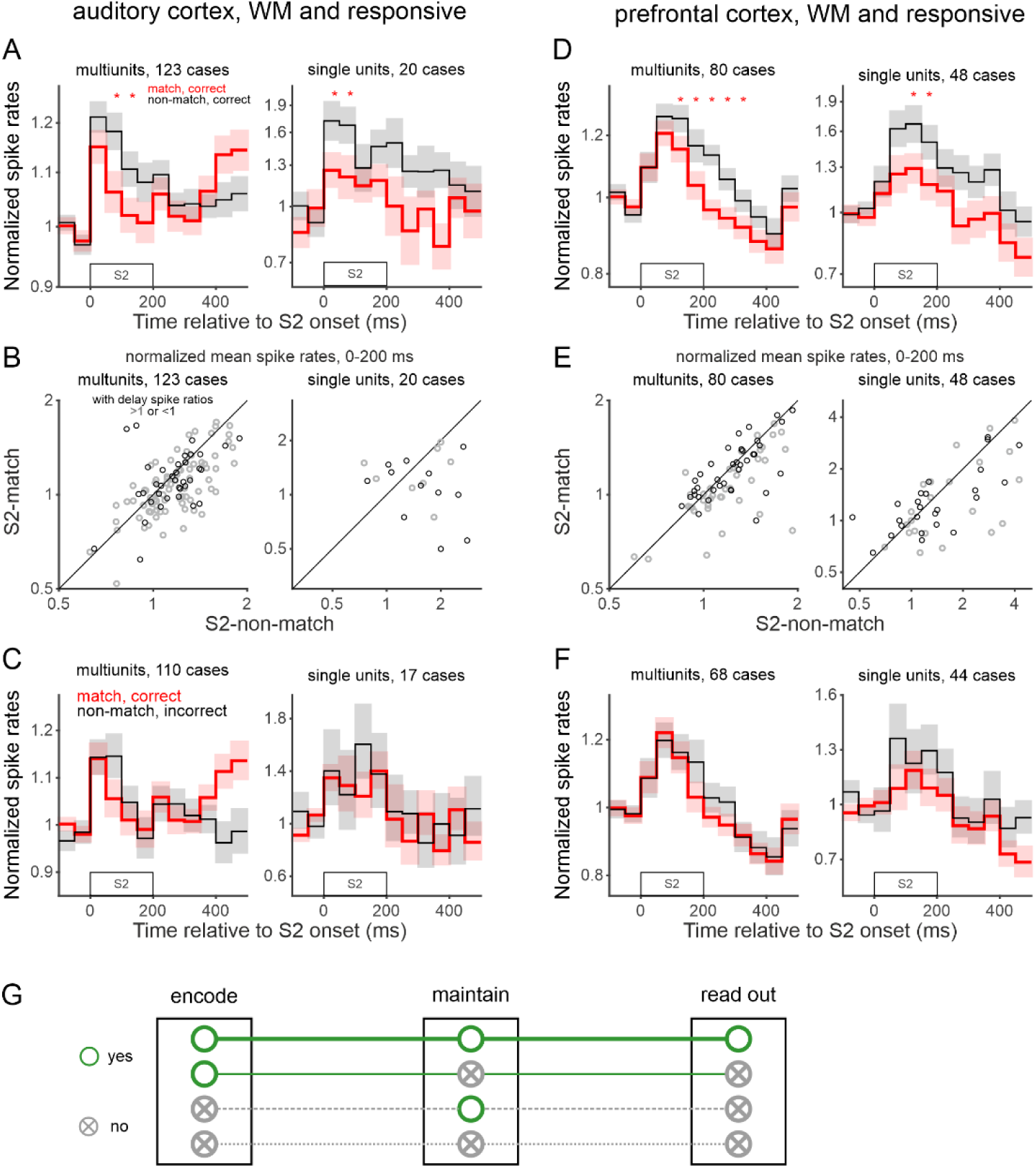
Working memory units in auditory and prefrontal cortices are also engaged in the readout of maintained information to guide behavior. (A-C) Readout activity of tone-responsive WM units in AC. Data were taken from cases where MUs and SUs responded to the 0.4-, 1.2-, or 3.6-kHz tone and exhibited significant delay spike ratios for the corresponding tone. (A) Population spike rates to S2 during correct task performance, computed with 50-ms bins, were generally weaker when S2 matched S1 (red) than when it did not (black). Asterisks mark significant differences between these two conditions. Note that significant differences were confined to the 200-ms S2 presentation window and disappeared hundreds of milliseconds before the monkeys released the bar in match trials (median reaction time of 624 ms relative to S2 offset). (B) These results were confirmed by comparing the average spike rates of individual units from 0 to 200 ms after S2 onset between the two conditions. Gray and black dots mark unit cases with significant delay spike ratios >1 and <1, respectively. (C) Population spike rates relative to S2 did not differ significantly between correct match trials and incorrect non-match trials in which the monkeys also released the bar. These trials types, however, differed in forward inhibition induced by S1. **(D-F)** Readout activity of tone-responsive WM units in PFC. Conventions are identical to those for panels A-C. **(G)** Schematic illustrating the four classes of neuronal populations in AC and PFC involved across the three processes required for performing a WM task: encoding, maintaining and reading out information. Open circles indicate population involvement in a given process, whereas circles containing a cross indicate no involvement. Horizontal lines connecting the symbols represent the same neuronal population across processes. Only the population that both encode and maintain information participate in its subsequent readout to guide behavioral responses.

Contributions of motor execution were excluded based on two findings. First, spike rate differences between match and non-match conditions decreased as bar release in match trials approached (median reaction time [RT] = 624 ms relative to S2 offset; 99.0% of RTs > 250 ms; Fig. S1A), becoming non-significant after S2 offset for both MUs and SUs (Fig. 3A). Second, bar release was associated with a spike rate increase rather than a decrease, as revealed by analyzing the release-aligned activity across all 330 MUs and 200 SUs (Fig. S1B). This increase began approximately 250 ms before the release, thus not overlapping with S2 presentation in 99.0% of trials. These findings rule out bar release as the cause of the match-related effects; otherwise, they would have manifested primarily as an elevation rather than a reduction and intensified as the release approached.

To estimate the contributions of forward inhibition, we compared population spike rates between correct match trials and incorrect non-match trials, the two conditions differing in this factor but both involving a Go response. Spike rates did not significantly differ between conditions during or after S2 presentation for either MUs or SUs (Fig. 3C; red vs. black traces). Therefore, forward inhibition could not explain the spike rate differences between correct match and correct non-match conditions.

Based on these results, we conclude that tone-responsive WM units were also involved in reading out information to guide behavior. They are in contrast to WM units that were not tone-responsive, whose population spike rates after S2 onset did not differ significantly between correct match and non-match conditions (Fig. S2A). These findings suggest a functional distinction between stimulus-encoding and non-encoding WM neurons during readout, despite both maintaining information during the delay.

Furthermore, tone-responsive WM units differed in their readout involvement also from units that did not maintain information during the delay, namely, those not exhibiting significant delay spike ratios for any of the three tones. These non-WM units showed no significant differences in spike rates between match and non-match conditions during S2 presentation, irrespective of whether they were tone responsive (Fig. S2B) or not (Fig. S2C). Therefore, these non-WM units either did not participate in readout or contributed via different mechanisms. After S2, spike rates became slightly higher in the match condition, reaching statistical significance for tone-responsive MUs (Fig. S2B, left); this possibly reflected the motor execution of bar release in these trials.

Taken together, these results indicate a selective role of stimulus-encoding WM neurons in reading out maintained information to guide actions. These neurons form a core network that bridges information encoding, maintenance, and readout to support goal-directed behavior (Fig. 3G, first row). They differed markedly from other populations, including stimulus-encoding neurons that did not maintain information during the delay (second row) and non-encoding neurons that either did or did not maintain information (third and fourth rows), none of which showed evidence of participating in readout according to our classification.

### 2.3. Working memory units in auditory cortex differ from units showing delay-baseline modulation

Isolating the mnemonic component of delay activity using our dual-task paradigm allowed us to identify genuine WM units, revealing a selective subset involved in both maintenance and readout. To determine how these WM units relate to those classified through the traditional single-task approach of comparing delay and baseline in the DMS task, we examined the overlap between these two populations. For this traditional metric, we calculated the DMS spike ratio, defined as the mean spike rate during the final 500 ms of the delay divided by that during the 500-ms baseline, separately for sequences with a 0.4-, 1.2-, or 3.6-kHz S1.

Strikingly, in nearly half of WM MUs and in about 75% of WM SUs (identified by the dual-task paradigm), DMS spike ratios remained close to 1 and were non-significant (p>0.05, permutation test; open dots in Figs. 4A, S3A, and S4A for the three sequences, respectively; for an example, see Fig. 1B, top-right panel). Therefore, a substantial proportion of WM units would remain undetected using the single-task approach.

**Fig. 4.**
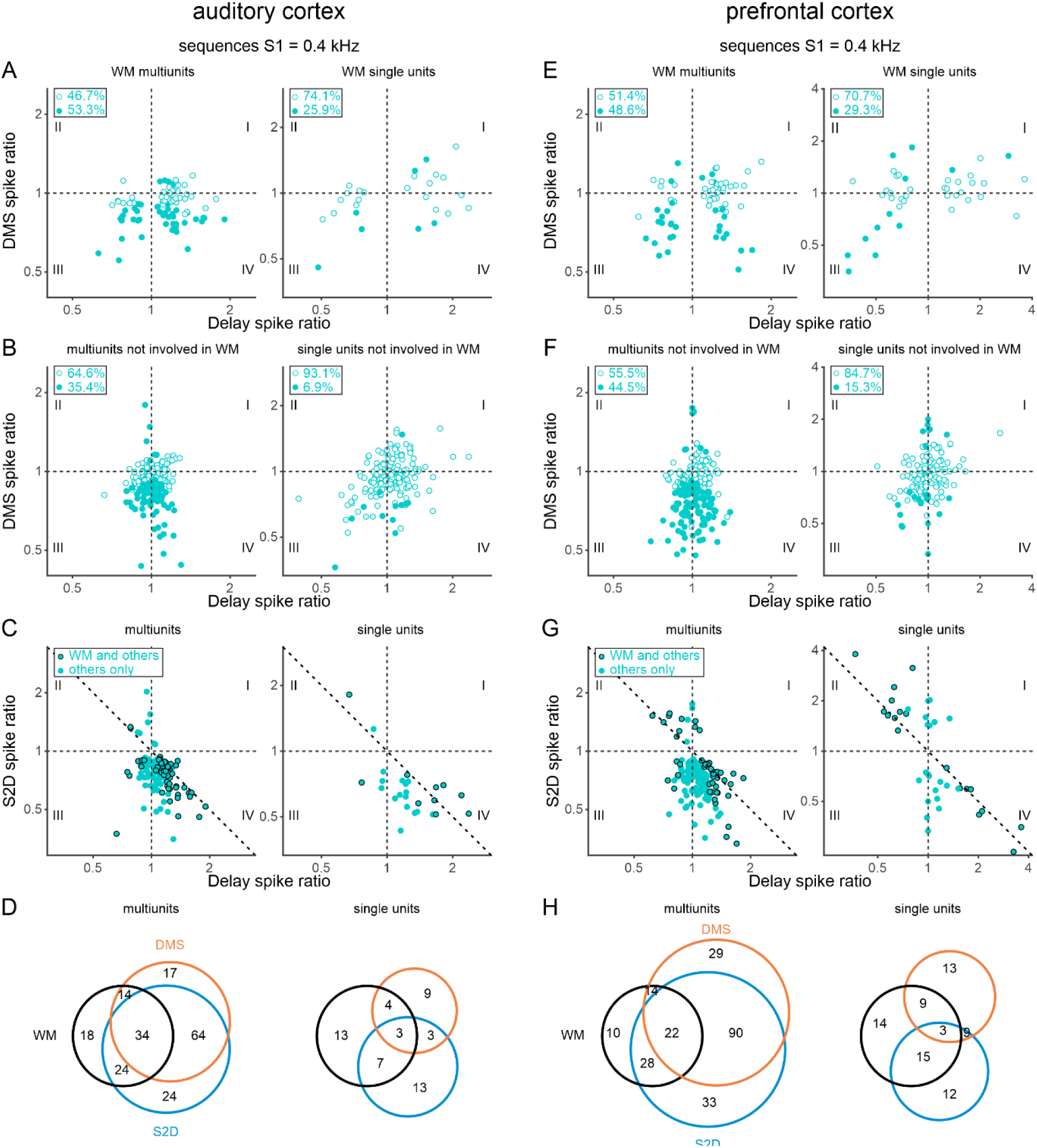
Limitations of the single-task approach for identifying working memory-related delay activity in auditory and prefrontal cortices. The single-task approach compares delay and baseline periods within the DMS task, expressed as DMS spike ratio. Data were obtained from correct trials with a 0.4-kHz S1. (A) Relationship between delay spike ratios and DMS spike ratios in AC WM MUs (left) and SUs (right). Filled and open dots represent units with and without significant DMS spike ratios, respectively. Only about half of the WM MUs and one-third of the WM SUs had significant DMS spike ratios, reflecting significant delay-baseline activity changes, and would also be identified as WM units using the single-task approach. The remaining large fraction of WM units were missed by the single-task approach. **(B)** The same relationship in AC units without WM-related delay activity. A substantial proportion of these units exhibited significant DMS spike ratios. **(C)** Effects of task-related mental processes other than auditory WM on spiking activity in AC. These effects were quantified by significant delay-baseline changes in the S2D task, expressed as S2D spike ratios. Dots with and without black edges mark units with and without WM-related delay activity, respectively. Most WM units showed significant delay spike ratios >1 but S2D spike ratios <1 (quadrant IV), or vice versa (quadrant II), demonstrating opposing effects of auditory WM and other mental processes on their spiking activity. **(D)** Venn diagrams showing the intersections among AC WM units (black), units with significant delay-baseline changes in the DMS task (orange), and units with delay-baseline changes in the S2D task (blue). The exact number of units within each section is provided. **(E-H)** Similar limitations of the single-task approach in PFC. Conventions are identical to those for panels A-D, evaluated for PFC MUs and SUs.

The remaining WM units had significant DMS spike ratios, making them detectable via the single-task approach (filled dots; Figs. 4A, S3A, S4A). Most of these significant ratios were <1, indicating that these units predominantly had suppressed spike rates during the delay compared to baseline. However, this ‘delay suppression’ failed to capture the genuine WM-related activity in many units, which instead increased when WM was required (filled dots, quadrant IV). A similar limitation applied to WM units showing ‘delay enhancement’ (significant DMS ratios >1): some units actually decreased spike rates when WM was required, resulting in delay spike ratios <1 (filled dots, quadrant II). Overall, such directional dissociations occurred in roughly half of the MUs (52.3% - 55.6% across sequences) and one-third of the SUs (31.0% - 37.0%). The single-task approach can thus obscure the directional nature of WM-related modulation.

Findings from units that were not identified as WM units by the dual-task paradigm further highlighted the discrepancy between approaches (Figs. 4B, S3B, S4B). About 40% of these MUs and 10% of these SUs nonetheless had significant DMS spike ratios (filled dots) and would thus be categorized as WM units using the single-task approach, despite exhibiting similar spike rates during the delay across the two tasks regardless of WM demand. Like the WM units, most of these ratios were <1, indicating that ‘delay suppression’ predominated even in units not involved in WM (filled dots in quadrant III and IV).

The discrepancy became even more apparent when directly comparing the distribution of WM units to units showing significant delay-baseline changes in the DMS task (‘DMS units’; Figs. 4D, S3D, S4D). The two populations diverged substantially: up to 81.3% of WM units showed no delay-baseline changes (Fig. S4D right; black vs. orange circles), while as many as 75.0% of DMS units were not involved in WM (Fig. S3D right). These results clarify that comparing the memory delay and baseline within a single task is often not sufficient to isolate WM-related activity.

Why did WM units and DMS units diverge? In the DMS task, delay and baseline periods differed not only in auditory WM but also in mental processes such as anticipation, decision making, and action preparation. Comparing delay to baseline in the S2D task, which differed only in processes unrelated to auditory WM, revealed that over 35% of 330 MUs and about 14% of 200 SUs showed different spike rates between these two periods, yielding significant S2D spike ratios (‘S2D units’; dots in Figs. 4C, S3C, S4C). Most of these ratios were <1 (93.5% - 93.8% of MUs and 82.1% - 92.3% of SUs across sequences), indicating that processes other than auditory WM can affect spiking activity, predominantly in a suppressive manner. These effects were observed in many units not involved in WM (dots without black edges; blue vs. black circles in Figs. 4D, S3D, S4D) and would cause delay-baseline changes in the DMS task, explaining why some DMS units are not genuine WM units.

These other mental processes could also affect spiking activity simultaneously with auditory WM, as reflected by the partial overlap between S2D units and WM units (dots with black edges in Figs. 4C, S3C, S4C; blue vs. black circles in Figs. 4D, S3D, S4D). Comprising up to 64.4% of WM MUs and 37.0% of WM SUs, this overlap allowed direct comparison between the effects of other processes (S2D spike ratio) and those of auditory WM (delay spike ratio). Notably, over 80% of these units had a significant S2D spike ratio <1 but a delay spike ratio >1 (Figs. 4C, S3C, S4C; quadrant IV), or vice versa (quadrant II). This indicates that auditory WM and other processes typically modulated spiking activity in opposite directions. In some cases, effects of other processes were even stronger, with S2D spike ratios deviating further from 1 than delay spike ratios (dots above the diagonal in quadrant II and below it in quadrant IV). These opposing influences explain why many WM units either showed no delay-baseline changes in the DMS task or exhibited changes that masked genuine WM activity.

### 2.4. Prefrontal cortex mirrors auditory cortex during working memory maintenance and readout

Is the maintenance-readout link at the level of individual units specific to AC or shared by higher-order executive regions? To address this question, we extended our analysis to PFC, an executive region widely considered crucial for WM. We applied the identical analytical pipeline to 346 MUs and 185 SUs recorded from the left vlPFC in 115 non-pharmacological sessions.

PFC results mirrored those from AC. During the delay, numerous PFC units exhibited different spike rates between the DMS and S2D tasks after certain S1s (see Fig. 1D for example units), reflecting their involvement in maintaining auditory information (Fig. 1E for analysis of trials with a 0.4-kHz S1). These WM units (141 of 346 MUs; 77 of 185 SUs) were distributed across both tone-responsive and non-responsive sub-populations (Fig. 2E). They exhibited tone-specific WM-related activity (Fig. 2F), persisting throughout the delay and ramping to a peak immediately before S2 (Figs. 2G, 2H). Moreover, during the subsequent readout phase, the tone-responsive subset of the WM units, but not the non-responsive subset or the non-WM units, differentiated match from non-match trials, exhibiting a match-related spiking reduction that reflected the monkeys’ match judgements and Go decisions (Figs. 3D-G, S2D-F, S1C). Finally, these genuine WM units diverged from the units identified using the single-task approach (Figs. 4E-H, S3E-H, S4E-H). Therefore, our dual-task paradigm reveals a functional principle shared by auditory and prefrontal cortices: integrating memory maintenance and readout within the same neurons.

### 2.5. Auditory cortex activity is necessary for working memory performance

While our electrophysiological findings demonstrate that neurons in AC are involved in WM, these findings remain correlational. To test whether AC activity is necessary for WM performance, we conducted additional perturbation sessions in Monkey L, microinjecting the dopamine D1 antagonist SCH23390 into the right AC, the same brain region where the data described above were obtained. In each session, we injected approximately 3.5 µL of saline (vehicle) or SCH23390 (7.5-40 mM concentrations) immediately before task performance. We analyzed behavioral accuracy (percentage of correct trials) across 42 sessions in which the monkey completed at least 20 trials per task.

The perturbation reduced behavioral accuracy in the DMS task in a concentration-dependent manner (Fig. 5A, left): with accuracy being generally lower at high concentrations (30 and 40 mM) than low concentrations (vehicle, 7.5 and 15 mM). A Kruskal-Wallis test confirmed a significant main effect of perturbation (p<0.05), with both high concentrations differing significantly from all three low concentrations (post-hoc Wilcoxon rank-sum tests; α = 0.005, corrected across ten comparisons).

**Fig. 5.**
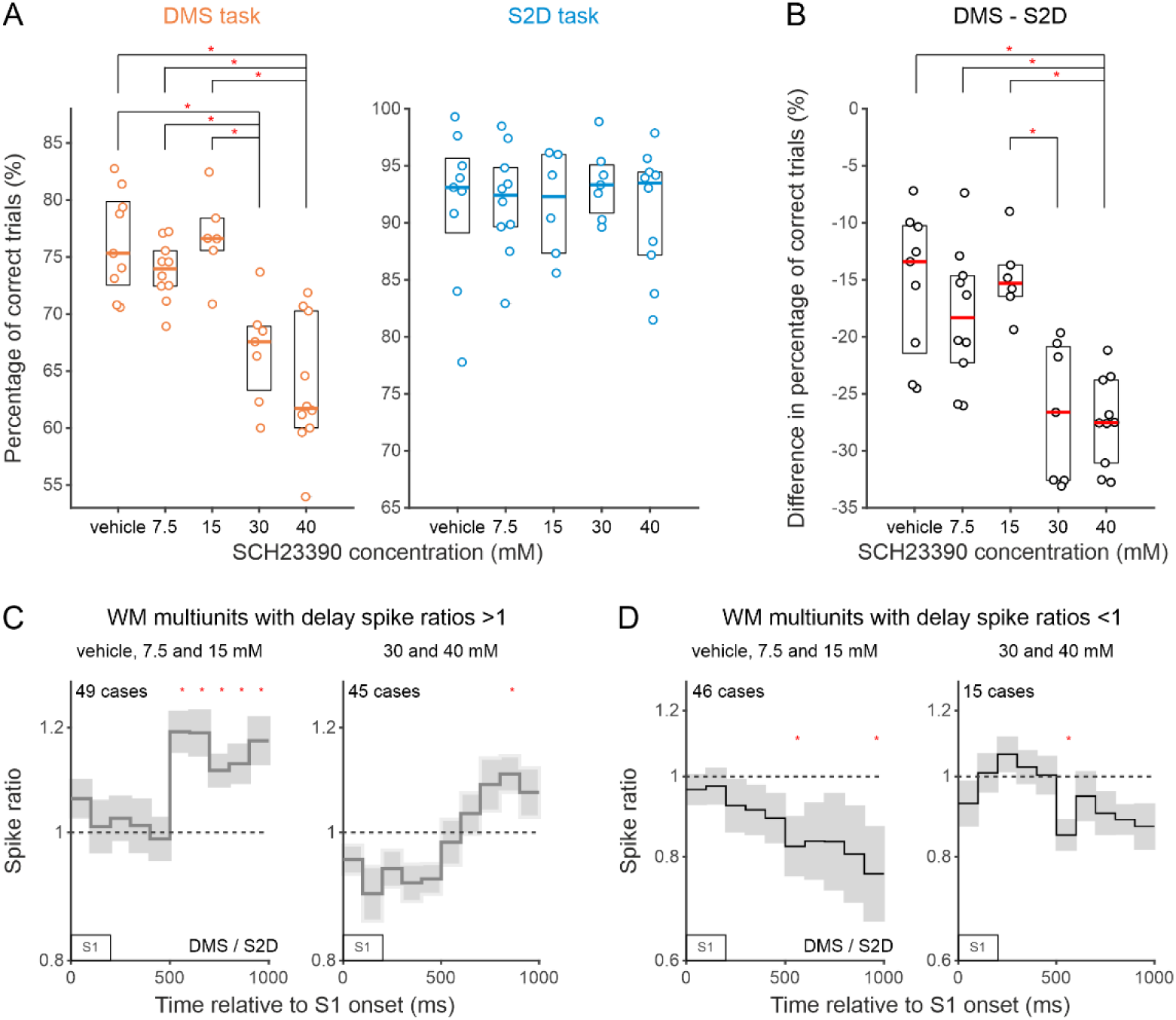
D1 receptor blockade in auditory cortex impairs working memory performance and memory-related delay activity. **(A)** Perturbation of AC activity via microinjections of the D1 receptor antagonist SCH23390 can reduce the percentage of correct trials in the DMS task (left) but not the S2D task (right). In each concentration condition, behavioral data for both tasks were collected within the same sessions. Dots represent individual session values. Boxplots illustrate the median (horizontal line) and interquartile range (box) across sessions. Asterisks denote significant differences between conditions. **(B)** Within-session differences in the percentage of correct trials between tasks were more pronounced at 30 and 40 mM concentrations. **(C, D)** Blocking D1 receptors in AC attenuates the differentiation between the DMS and S2D tasks during the delay. Population spike ratios between the two tasks are plotted relative to S1 onset for WM MUs identified under low (left; vehicle, 7.5, and 15 mM) and high concentrations (right; 30 and 40 mM), shown separately for MU cases with significant delay spike ratios > 1 (C) or < 1 (D). Deviations of the population spike ratios from 1 reflect the degree of between-task differentiation. Note that this task differentiation during the delay was markedly reduced and disrupted under high concentrations relative to that under low concentrations.

The reduced accuracy in the DMS task was selectively driven by the impairment of auditory WM rather than other mental processes such as stimulus and reward perception, anticipation, decision making, or motor execution. Accordingly, the perturbation did not significantly alter behavioral accuracy in the S2D task, which shared all processes with the DMS task except auditory WM (Fig. 5A, right; Kruskal-Wallis test, p>0.05). Further support was obtained by assessing within-session accuracy differences between tasks. These differences were significantly modulated by the perturbation, widening at high concentrations (Fig. 5B; Kruskal-Wallis main effect, *p* < 0.05). Post-hoc comparisons showed that 40 mM differed significantly from all three low concentrations, while 30 mM differed significantly from 15 mM (ten Wilcoxon rank-sum tests, α = 0.005). Therefore, local perturbation of AC D1 receptors specifically impairs auditory WM, demonstrating a causal link between AC activity and WM performance.

To determine how this behavioral degradation relates to underlying neural changes, we analyzed 111 WM MUs across the 42 perturbation sessions (31.2% of 356 recorded MUs). These WM MUs were identified using the same criteria established in Section 2.2 (exhibiting significant delay spike ratios >1 or < 1 in correct trials after the 0.4-, 1.2-, or 3.6-kHz S1s), reaffirming the consistent prevalence of WM-related activity in AC. To evaluate local populations’ overall differentiation between tasks across drug concentrations, we pooled correct and incorrect trials for the subsequent analysis. This avoids biasing the analysis toward trials where the animal successfully compensated for the perturbation or failed due to transient behavioral fluctuations. Spike ratios between tasks were calculated in 100-ms bins aligned to S1 onset for each unit, restricted to cases with significant delay spike ratios after specific S1 tones during WM unit identification.

For cases with significant delay spike ratios >1 (Fig. 5C), population spike ratios under low concentrations (left; vehicle, 7.5 and 15 mM) deviated significantly from 1 during the 1000 ms after S1 onset, with significance persisting across all five bins in the final 500 ms of the delay (permutation test, p<0.05/10, corrected across ten bins). Under high concentrations (right; 30 and 40 mM), this task differentiation was markedly attenuated and temporally truncated, with the profiles shifting closer to 1 and significance restricted to a single bin. This drug-induced degradation was also observed for cases with significant delay spike ratios <1 (Fig. 5D): the population spike ratios showed greater deviation from 1 under low than under high concentrations (left vs. right). Together, these results demonstrate that blocking D1 receptors in AC attenuates the local population’s capacity to maintain task-related auditory information, establishing the necessity of AC for accurate WM performance.

## 3. Discussion

This study provides electrophysiological evidence that neurons in AC and PFC exhibit spiking activity associated with the temporary maintenance of auditory information, and uncovers that a distinct subset of these WM neurons, specifically those that also encode the auditory stimulus, participates directly in the readout of maintained information to guide subsequent behavior. These findings bridge the two key phases of WM - maintenance and readout - within the same neurons, offering a unified framework for understanding the neural implementation of WM across the sensory-executive axis. Furthermore, our perturbation results in one monkey establish the causal necessity of the sensory node within this axis, as manipulating AC activity impaired WM performance. Crucially, these insights relied on a dual-task paradigm designed to dissociate WM from other concurrent mental processes and task variables. With this approach, we identified WM neurons that only partially overlapped with the memory neurons traditionally defined by delay-baseline comparisons within a single task. This indicates that single-task approaches provide an incomplete profile of WM circuits. Our findings thus prompt a re-examination of prior efforts to localize WM neurons and model the underlying neuronal mechanisms, not only in the auditory domain but also in other sensory modalities.

Our finding of WM neurons in both AC and PFC provides electrophysiological support for the distributed network models, which, however, are based mainly on visual fMRI studies, positing that WM recruits a broad cortical network spanning from sensory areas to higher-order executive regions^2,12,33,35,37^. Beyond this macro-level framework, our data extend these models by revealing a specialized population of neurons in both AC and PFC that directly bridges maintenance and readout during WM tasks. This within-circuit continuity stands in contrast with the weak coupling observed between neuronal signals during the memory delay and the readout phase in previous studies^26–32^, highlighting a fundamental principle of the cellular implementation of WM across the sensory-executive axis. Ultimately, our findings help reconcile the ongoing debate over whether memory maintenance and readout are anatomically segregated or functionally integrated^2,11,33–37^.

In AC and PFC, the neurons bridging maintenance and readout are predominantly confined to the population that responds to the to-be-remembered tone during its presentation, rather than the non-responsive population. Neurons within this latter group remain silent during tone presentation yet still exhibit WM-related delay activity, indicating that they specialize in maintaining information in the absence of direct sensory drive. They may maintain primarily information that is more abstract, such as learned associations of tones with actions. In contrast, the responsive neurons may maintain both sensory features, like frequency, and abstract associations. This functional distinction in maintenance may underlie their differential involvement in readout, where sensory features of S1 must be retrieved and compared with those of S2. Therefore, a neuron’s response profile during stimulus processing may not only shape its role in maintaining information during the delay, but also its ultimate capacity to translate mnemonic representations into appropriate actions according to task demands.

The essential role of these WM neurons in AC for behavioral performance is supported by our perturbation results. Attenuating the mnemonic component of delay activity in these neurons, via blocking D1 receptors within AC, was accompanied by an impairment in WM performance. These results establish the necessity of AC for WM, a finding in line with the observations that lesions or inactivation of AC impair performance on memory tasks^51–54^. Our findings thus expand the framework of the sensory-executive axis in WM: the sensory node is a critical anchor in its own right, rather than a passive recipient of frontal top-down signals. Our findings further extend the known importance of the dopaminergic system in WM from PFC to sensory cortex^46,47^, suggesting that dopamine is a key neuromodulator underlying WM across this functional axis.

A subset of the AC and PFC WM neurons identified via our dual-task paradigm displayed delay-baseline changes in the DMS task. This suggests that, in previous auditory WM studies using the single-task approach, some of the AC or PFC neurons showing delay enhancement or suppression relative to baseline might be genuinely involved in the memory process^29–32,45,54^. Going beyond these traditional delay-baseline comparisons, we characterized WM-related delay activity as spike rate differences during the delay between two tasks with distinct WM demands. In most AC and PFC neurons, this activity was present throughout the entire delay period, supporting WM models that rely on persistent activity^1,3,55^. Crucially, by showing that local D1 receptor blockade in AC attenuates this persistent signal alongside impairments in WM performance, our results provide direct evidence for its causal necessity in WM. In addition, this WM-related activity often became more prominent toward the end of the delay shortly prior to S2 presentation. Such ramping dynamics are consistent with the framework that slow firing changes in AC may provide a neuronal mechanism for memorizing and associating behaviorally relevant events^19^. By isolating the memory-specific component of these dynamics from other task-related variables, our results lend weight to the idea that AC utilizes a tonic representational system to bridge events across time, complementing the transient, stimulus-driven representations.

The ramping dynamics of the WM-related delay activity suggest that, in addition to features of the past S1, this activity also carries information about future tones upcoming at S2, including their associated behavioral responses. This is consistent with the future-oriented function of WM in transforming past sensory information into memory-guided actions, a framework suggested by studies in the visual modality^36,56,57^. Such a proactive capacity is further supported by our finding that only units exhibiting this delay activity fired in a manner linked to the monkeys’ match judgements and behavioral decisions during the readout phase. Activity related to these cognitive processes has been observed in AC and PFC during auditory tasks^32,58–60^, and it aligns with our previous findings that neuronal responses in these regions convey the behavioral meaning of sounds^43,44^. Consistent with Hwang and Romanski^32^, spike rates during S2 presentation were generally lower when S2 matched S1 than when it did not. Although this phenomenon is often termed ‘match suppression’, it may result from a suppression of activity in the match condition, an enhancement in the non-match condition, or both. The predominance of ‘match suppression’ in our study contrasts with reports of comparable suppression and enhancement in other WM studies^26–31,45^. This discrepancy may arise because the match effects observed in those paradigms could reflect not only judgements or behavioral decisions, but also differences in forward inhibition between match and non-match trials^50,61^ or activity directly associated with motor execution.

In summary, by strictly isolating auditory WM from other concurrent task-related factors, our study identifies a fundamental principle of its cellular implementation: memory maintenance and behavioral readout converge within individual neurons. This convergent cellular architecture is conserved across the sensory-executive axis. However, it remains a critical open question whether this functional organization generalizes to WM in other sensory modalities. Non-WM processes, such as stimulus and reward anticipation, decision making, and action preparation, exert profound influences across visual, somatosensory, and olfactory areas, as well as higher-order regions like parietal cortex^23,62–68^. Disentangling these concurrent variables remains a major challenge, particularly in single-task paradigms where delay-baseline modulations may multiplex task factors. As neuronal mechanisms of auditory memory could well differ from those in other modalities^53^, it is possible that the convergent cellular WM architecture is unique to the auditory system rather than a universal cross-modal principle.

## 4. STAR methods

### 4.1. Experimental model and subjects

Monkey C and Monkey L (Macaca fascicularis) participated in this study. They were implanted with a head holder to allow atraumatic head fixation and two recording chambers to allow access to cortical regions with microelectrodes (for details of the implantation, see Brosch and Scheich^61^). One chamber was implanted over the right AC and the other over the left PFC. The two animals were euthanized with an overdose of Nembutal after completing either all non-pharmacological recording sessions or all perturbation sessions. The authority for animal care and ethics of the federal state of Saxony-Anhalt approved this study, which conformed to the rules for animal experimentation of the European Communities Council Directive (86/609/EEC).

### 4.2. Behavioral paradigms

Both monkeys were trained to perform both the delayed match-to-sample (DMS) task and the S2-discrimination (S2D) task. In these tasks, a trial started by illuminating LED lights. Within the next 4 s, the monkeys had to grasp a touch bar and hold onto it. After maintaining the holding for ∼1.5 s, a sequence of two sounds, S1 and S2, was presented. The sounds had a duration of 200 ms and they were separated by a silent delay of 800 ms. Within a specific time window after S2 offset (40-1760 ms), the monkeys had to make either a *Go* response (i.e., release the bar) or a *NoGo* response (i.e., continue holding the bar throughout the window) to earn a fluid reward. Which response was required in a trial depended on tasks and sound sequences. In trials with a correct response, the lights were turned off when the reward was delivered. In incorrect trials, the lights were turned off when the monkeys released the bar or when they did not grasp the bar within 4 s after light onset. The lights were turned off for at least 5 s.

In both tasks, S1 could be a 0.4-, 1.2-, or 3.6-kHz tone. These tones were also used as S2 in the DMS task. In this task, a Go response was required when S1 and S2 were the same and a NoGo response when they were different. In contrast, S2 in the S2D task could be a noise burst, a 20-Hz click train, or an 80-Hz click train. In this task, a Go response was required when S2 was the noise burst and a NoGo response when it was a 20-Hz or 80-Hz click train. All sounds were presented bilaterally through free-field loudspeakers (Karat 720, Canton, Weilrod, Germany) and had a sound pressure level of approximately 67 dB SPL, measured with a free-field ½ inch microphone (40AC, G.R.A.S., Vedbaek, Denmark) placed near the center of the monkeys’ head.

The two tasks were performed in separate, alternating blocks of about 140 trials in each experimental session without pharmacological perturbation. The block size was reduced to about 30 trials in sessions with the perturbation to ensure that both tasks were completed within a comparable temporal window of drug efficacy. Go trials constituted about 50% of the trials in each block. The order of blocks and the starting task counterbalanced across sessions. The two tasks were distinguished by LED lights. Red LEDs located on the left side of the monkeys were used in the DMS task, whereas green LEDs located on the right side of the monkeys were used in the S2D task (for further details, see Huang and Brosch^25^).

In this report, we utilized neuronal data obtained from 134 experimental sessions without pharmacological perturbation (72 from Monkey C and 62 from Monkey L). In each of these sessions, the monkeys correctly performed both the DMS and S2D task above the 50% chance level (permutation test, p<0.05). Across these sessions, the percentage of correct trials ranged from 60% to 80% (median of 70%) in the DMS task and from 67% to 96% (median of 87%) in the S2D task.

Following the completion of these non-pharmacological sessions, we incorporated behavioral and neuronal data from an additional 42 sessions with pharmacological perturbation in Monkey L. During these sessions, the monkey performed the two tasks immediately after receiving intracortical injections into AC. These injections consisted of either saline (vehicle) or the D1 antagonist SCH23390 (Sigma-Aldrich Chemie GmbH, Steinheim, Germany) dissolved in saline at one of four concentrations: 7.5, 15, 30, or 40 mM. The total volume delivered in each session ranged from 3.0 to 4.5 µL, administered at an injection rate of approximately 0.3 µL/minute (range: 0.2-0.6). Median volumes for the five conditions (vehicle through 40 mM) were 3.4, 3.6, 3.6, 3.5, and 3.5 µL, respectively. The five conditions were tested in blocks of five consecutive sessions, with their order randomized within each five-day block. The total number of sessions included in the analysis for the five conditions was 9, 10, 6, 7, and 10, respectively.

### 4.3. Passive paradigms

In the passive conditions, the monkeys were presented with the same set of sound sequences as in the two tasks. Here, the LEDs were not illuminated, the touch bar was not functional, and no water was delivered. The nine sequences used in the DMS task were presented in half of the blocks, while the nine sequences used in the S2D task were presented in the other blocks. These passive conditions were conducted after the monkeys had performed the tasks in 90 of the 134 non-pharmacological sessions, as well as in all 42 perturbation sessions (though passive data under perturbation were not included in the present study).

Before or after the monkeys performed the tasks, we also presented 400 pure tones with 40 different frequencies, typically covering a range of eight octaves in equal logarithmic steps, to measure the best frequency and the frequency response area of each unit^61^. In addition, we presented complex sounds, such as noise bursts, dog barking, monkey coos and human vocalizations, to assess the sound selectivity of the units^69^.

### 4.4. Electrophysiological recordings

Microelectrode recordings of neuronal activity started once the monkeys correctly performed both the DMS task and S2D tasks significantly above the 50% chance level during ten consecutive sessions (permutation test, p<0.05).

Recordings were conducted in the core fields of the right AC, mainly in field A1, and in the ventrolateral region of the left PFC (vlPFC). To determine the position of A1, we used the spatial distribution of the best frequencies across the recording sites and the information about the electrode tracks through the brain, e.g., whether the electrodes passed through parietal cortex into AC or not^70,71^. To identify the vlPFC, we used the characteristics of its responses to pure tones and complex sounds presented in the passive conditions (for details, see Huang and Brosch^69^) and the information about its anatomical location ventral to the principal sulcus and anterior to the arcuate sulcus^72,73^. Part of the data has been used to show that LED lights in the tasks can evoke neuronal responses in AC and PFC^25^.

During sessions without pharmacological perturbation, two multichannel manipulators (Thomas Recording, Giessen, Germany) equipped with up to seven microelectrodes (80-µm diameter) were used to record action potentials (0.5 - 5 kHz) simultaneously from AC and PFC. The action potentials of a few neurons (multiunit) recorded from a given microelectrode were isolated using the built-in spike detection tools of the data acquisition system (Alpha-Map, Alpha Omega, Grapeland, Israel). The time stamps and waveforms of the action potentials were stored. A principal component analysis was conducted offline on the waveforms of the action potentials to extract single-unit activity.

During sessions with pharmacological perturbation, an S-probe (236 µm diameter, 100 mm total length) with 16 single electrodes (15 µm diameter; 150 µm electrode spacing; 580 µm from the first electrode to the probe tip; Plexon, Dallas, USA) was used to record neuronal activity in AC via an Intan data acquisition system (Intan Technologies, Los Angeles, USA). Recorded broadband signals were band-pass filtered (0.5-5 kHz; 2^nd^-order Butterworth) and thresholded (mean ± 3 SD) to obtain multiunit activity.

### 4.5. Intra-cortical microinjections

Microinjections of saline or the D1 antagonist SCH23390 were performed through a customized fluid channel (640 µm diameter) located between electrodes 12 and 13 of the S-probe (Plexon, Texas, USA). Injection delivery was controlled by a pneumatic system (PMI-100 Pressure Micro-Injector; Dagan Corporation, Minneapolis, USA).

### 4.6. Data analyses

Data analyses were performed using MATLAB (MathWorks, Natick, MA, USA). To analyze task-related neuronal activity, we included only units responsive to sounds during the tasks. Similarly, for analyses in the passive conditions, only units responsive in these conditions were used.

To determine whether a unit was responsive to sounds during task performance, we computed twelve post-stimulus time histograms (PSTHs), each triggered to the onset of S1 or S2. These corresponded to the 0.4-, 1.2- and 3.6-kHz tones when they were presented in the two tasks, as well as the noise burst, the 20-Hz and the 80-Hz click train when they were presented at S2 in the S2D task. The PSTHs had a bin size of 10 ms and were computed by averaging spike rates across all trials in the conditions, irrespective of whether the monkeys performed the tasks correctly or not. A unit was considered responsive to sounds if, in any of the twelve PSTHs, spike rates in at least two consecutive bins from 0 to 200 ms after sound onset exceeded the baseline mean spike rate (calculated during the 100-ms period before sound onset) by more than three standard deviations (> mean + 3SD). Units were classified as non-responsive if their spike rates remained within three standard deviations of the baseline mean (mean ± 3SD) across all twelve PSTHs. Only these responsive and non-responsive units were included in the analysis. We also conducted this same classification procedure for the 0.4-kHz tone based on the three PSTHs triggered to this tone. The same logic was applied to the 1.2- and 3.6-kHz tones.

To determine whether a unit was responsive to sounds under the passive conditions, we performed corresponding analyses on the data obtained in these conditions.

To calculate spike ratios for a unit, we first computed for each condition a PSTH aligned to the onset of S1, using a bin size of 100 ms. The DMS spike ratio was defined as the mean spike rate during the final 500 ms of the delay divided by the mean firing rate during the 500-ms baseline preceding S1 onset in the DMS task. The S2D spike ratio was calculated in the same way, using the activity from the S2D task. Here, we used only units recorded in at least four trials, as this is the minimum number required to perform a permutation test with an alpha value of 0.05.

The mean spike rate during the final 500 ms of the delay in the DMS task was then divided by that in the S2D task to obtain the delay spike ratio. The baseline spike ratio was computed in the same way based on spike rates during the baseline. The delay spike ratios were also calculated between corresponding passive conditions.

Based on the PSTHs with a bin size of 100 ms, we computed population spike ratios between the two tasks. We first calculated for each unit the ratios of mean spike rates during consecutive 100-ms windows aligned to S1 onset, separately for sequences with a 0.4-, 1.2-, or 3.6-kHz S1. These ratios were normalized, that is, divided by the mean ratio during the 100 ms preceding S1, and then averaged across selected units and sequences. A population spike ratio in any of the ten windows after S1 onset was considered significant if p<0.005, reflecting a total alpha value of 0.05 across the ten windows (0.005×10).

To estimate the duration of persistent spike rate differences between the DMS task and S2D task, we computed PSTHs with a bin size of 50 ms. For each unit in trials with a given S1, we computed the ratio of the mean spike rate in the DMS task to that in the S2D task during 100-ms sliding windows, advanced in 50-ms steps, across the final 500 ms of the delay period. The duration of persistent spike rate differences was defined as the longest sequence of consecutive 100-ms windows across which the spike ratios remained consistent in direction relative to 1. This persistency duration was measured from the onset of the first window in the sequence to the offset of the last.

Population spike rates to S2 in a given condition were calculated from individual PSTHs with a 50-ms bin size and aligned to S2 onset. The PSTHs were divided by the mean spike rate during the 100 ms preceding S2 onset for normalization, and then averaged across selected units and sequences. Due to the small bin size, population spike rates during the 500 ms after S2 onset were considered significantly different between conditions if p<0.07 in at least two consecutive bins, a criterion that avoids spurious single-bin significance and corresponds approximately to a total alpha value of 0.05 across the ten bins (0.07×0.07×10).

The PSTHs aligned to S2 onset were also used to compute normalized mean spike rates during S2 presentation in a given condition, defined as the spike rate averaged over the 0-200 ms interval after S2 onset divided by the spike rate averaged over the 100 ms before S2.

Population spike rates to bar release were computed in the same manner as those to S2, based on PSTHs aligned to bar release and normalized to the 200-300 ms interval after bar release. Spike rates in each 50-ms bin from 300 ms before to 200 ms after bar release were compared with 1 and considered significant if p<0.07 in at least two consecutive bins.

In each pharmacological perturbation session, we quantified behavioural accuracy in the DMS and S2D tasks by calculating the percentage of correct trials, that is, the number of correct trials divided by the total number of trials the monkey performed. For this analysis, we included only sessions in which the monkey performed at least 20 trials in both tasks; this threshold ensured a minimum measurement resolution of 5%.

## Acknowledgements

This research was supported by the European Regional Development Fund (CBBS Neuronetwork 13) and the Deutsche Forschungsgemeinschaft (project number: 457346369).

