## Supplementa figures for "Individual neurons in monkey sensory and prefrontal cortices contribute to both working memory maintenance and readout"

**This PDF file includes:**

Figs. S1 to S4.

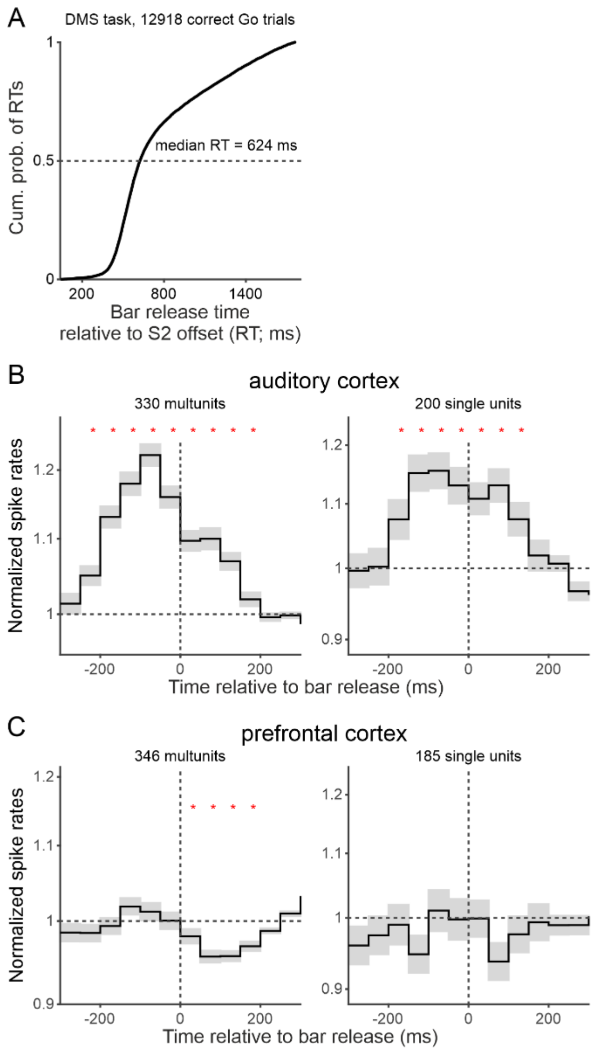


**Fig. S1. Spiking activity associated with bar release cannot account for the match-related effects during the readout phase in auditory and prefrontal cortices. (A)** Cumulative distribution of reaction times (RTs) for bar release across all 12,918 correct match trials performed by the two monkeys in 134 non-pharmacological sessions. RT was calculated as the difference between the time of bar release and the offset of S2. The median RT was 624 ms. **(B)** Population spike rates aligned to bar release across all 330 MUs (left) and 200 SUs (right) recorded from AC in the non-pharmacological sessions. Data were obtained from correct match trials in the DMS task. Stars mark 50-ms bins in which the population spike rates were significantly different from 1. Note that population spike rates were greater than 1 around bar release, reaching significance only from approximately 250 ms before to 200 ms after the release. (**C**) Population spike rates aligned to bar release across all 346 MUs (left) and 185 SUs (right) recorded from PFC in the non-pharmacological sessions. Conventions are identical to those of panel B.

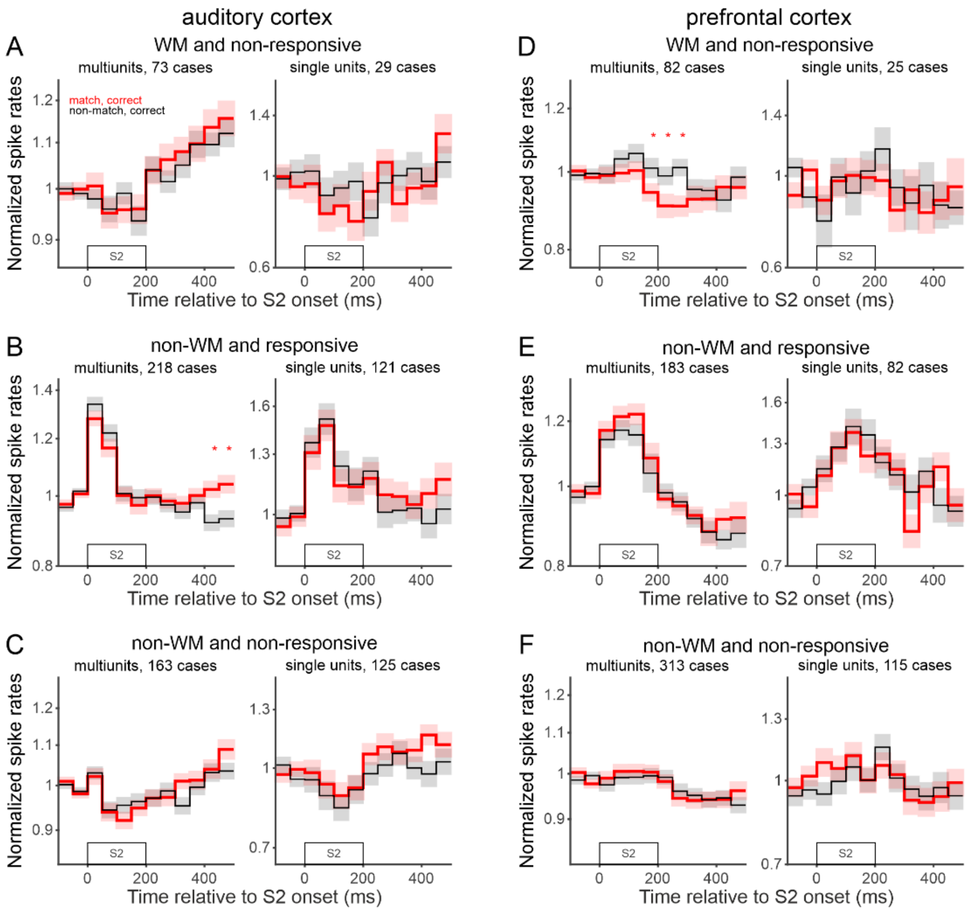


**Fig. S2. Readout activity is generally absent in non-responsive WM units and non-WM units in auditory and prefrontal cortices. (A)** Non-responsive WM units in AC exhibited no readout activity. Conventions same as those of Fig. 3A. **(B, C)** Non-WM units in AC exhibited no readout activity, irrespective of whether they were tone-responsive (B) or not (C). Non-WM units were those showing no significant delay spike ratios for any of the three tones. Conventions match those of panel A. **(D-F)** Readout activity was not consistently observed across non-responsive WM MUs (D, left) and SUs (D, right) in PFC, and was absent in non-WM units (E, F).

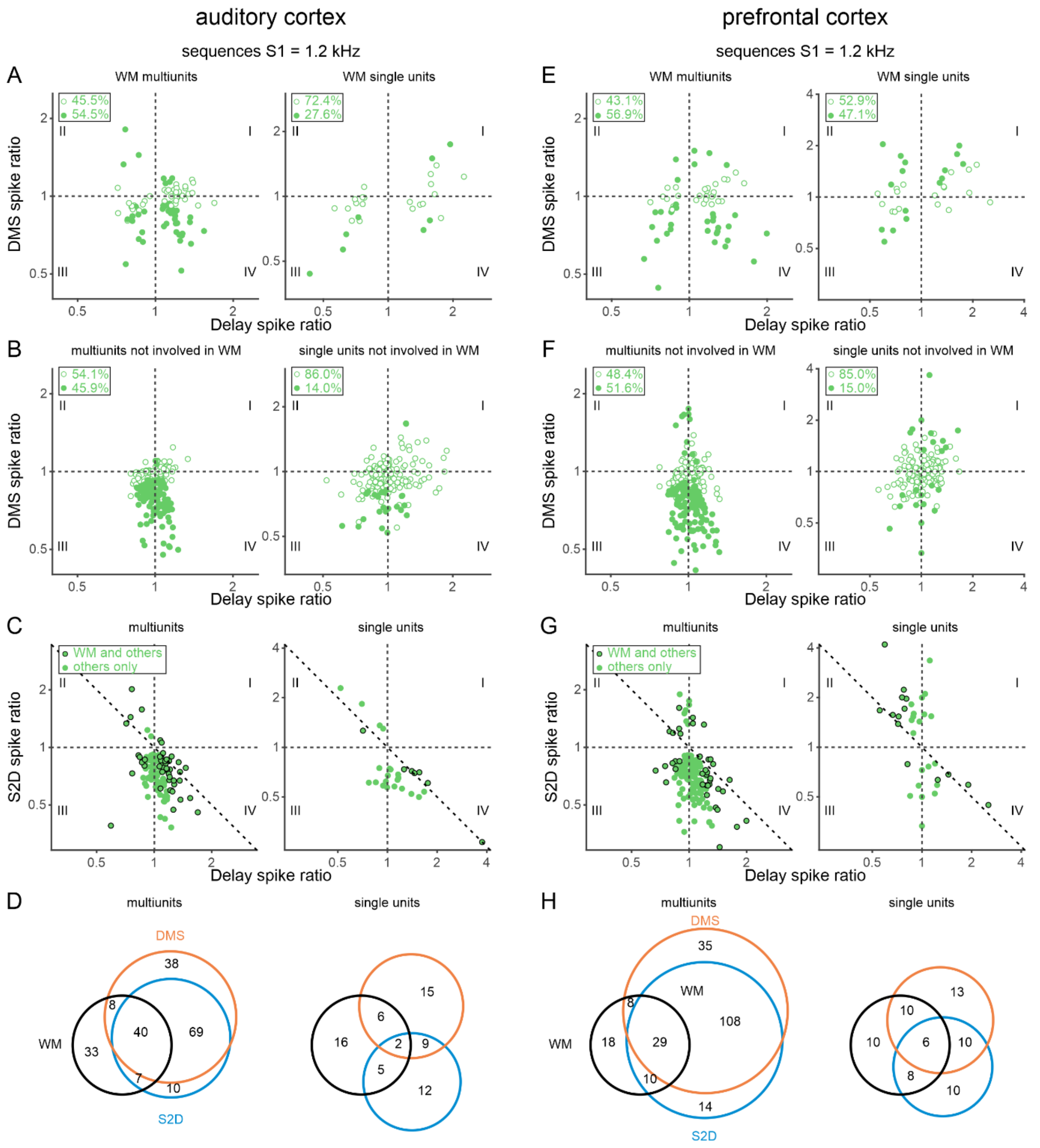


**Fig. S3. Limitations of the single-task approach for identifying working memory-related delay activity in auditory and prefrontal cortices: evidence from sound sequences with a 1.2-kH S1.** Conventions are as in Fig. 4.

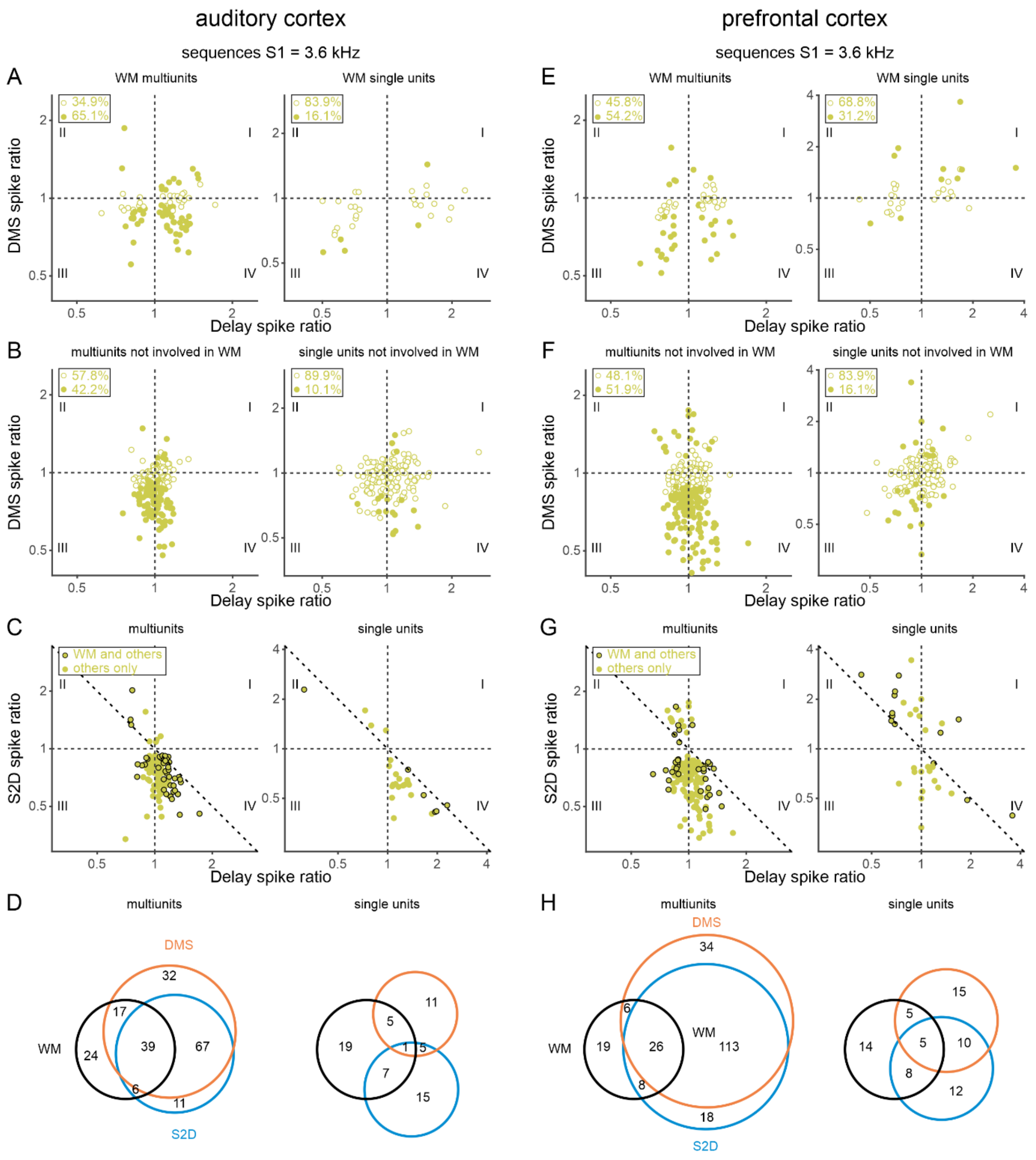


**Fig. S4. Limitations of the single-task approach for identifying working memory-related delay activity in auditory and prefrontal cortices: evidence from sound sequences with a 3.6-kH S1.** Conventions are as in Fig. 4.
